# Light-dependent and predator-inducible aldehyde synthesis in *Prochlorococcus* for specific defense against *Uronema*

**DOI:** 10.64898/2026.06.08.730813

**Authors:** Peichun Lin, Yelinzi Yuan, Xiaokun Liu, Zuyuan Gao, Tianzhuo Hou, Lingfeng Huang, Rui Zhang, Bi-Xian Mai, Zhili He, Hongbin Liu, Qinglu Zeng, Shanquan Wang

## Abstract

Cyanobacteria as a primary producer provide energy and carbon sources for the marine food web, of which predation-interactions play central roles in regulating global element cycles and marine ecosystem stability. Here, we report the anti-predation activity of a typical marine cyanobacterium *Prochlorococcus* MED4 to defend the predation by *Uronema marinum*. MED4 synthesize formaldehyde as the anti-predation chemical, of which the synthesis was light-dependent and predator-inducible. Compared to other protists, both the higher concentration of accumulated formaldehyde in *U. marinum* and the lower formaldehyde tolerance of *U. marinum* resulted in the specific anti-predation of MED4 against *U. marinum*. This specific anti-predation could regulate the cyanobacterial growth and the *U. marinum* infection of marine fishes. Metadata analyses showed the mutually exclusion of *Prochlorococcus* and *Uronema* in global marine environments. These findings significantly advance our understanding of the marine food web and biogeochemical cycles.

**Significance Statement:** Predation-driven interactions in the ocean are critical regulators of global biogeochemical cycles, yet active defense mechanisms in marine picophytoplankton remain largely unknown. This study reveals that *Prochlorococcus* MED4 as the most abundant primary producer in the ocean synthesize light-driven and predator-inducible aldehydes to actively and specifically defend predation against the ciliate *Uronema marinum*. The anti-predation of *Prochlorococcus* against *Uronema* could have broad implications in biocontrol of *Uronema* infection in marine fishes and regulation of the marine food web.

## Introduction

Marine cyanobacteria *Synechococcus* and *Prochlorococcus* are the most abundant photosynthetic organisms on Earth, numerically dominating most oceanic waters^1, 2^. Their core functions of oxygenic photosynthesis and carbon fixation play a pivotal role in shaping Earth’s early history and in mediating global biogeochemical cycles^3–6^. Based on their vast global populations with mean annual abundances of approximately 7.0 × 10^26^ cells for *Synechococcus* and 2.9 × 10^27^ cells for *Prochlorococcus*, they collectively contribute around 25% of the ocean net primary productivity^7^. Therefore, these picocyanobacteria are consequently believed to be among the most important organisms ever on the planet^6^. The *Prochlorococcus* adapt to oligotrophic oceans (particularly the euphotic zone between 50°N and 50°S) by reducing cell and genome sizes, which enhance nutrient uptake and light absorption efficiency, and lead to its global abundance surpassing that of *Synechococcus*^7–9^. The pivotal roles of *Prochlorococcus* in marine ecosystems enable that changes in its population dynamics can have far-reaching global impact on the ocean productivity, nutrient cycles and carbon sequestration processes. Though previous studies have suggested that temperature is a decisive environmental factor regulating the global distribution patterns of *Prochlorococcus* and *Synechococcus*^7, 10^, predation has been recently recognized as a key factor influencing the distribution and ecological dynamics of marine cyanobacteria^11, 12^.

The cyanobacteria as a primary producer through carbon sequestration provide the energy and carbon sources for microzooplankton (protists) and establish the microbial loop^13, 14^, in which the fixed carbon is transferred up the food chain, from primary consumers to mesozooplankton and ultimately to higher trophic levels, including fish and marine predators^15^. These cyanobacteria contribute to the global carbon cycle and the stability of marine ecosystems. Predation within this system is central to the energy transfer and element cycling by regulating population dynamics and maintaining ecological balance^16, 17^. The continuous predation pressure has been identified as a key driver of microbial evolution in terrestrial ecosystems, which promotes ecological adaptation and evolution in prey species and triggers the ongoing evolutionary arms race for developing strategies to counteract the predation pressure^18, 19^. Therefore, the anti-predation strategies may alter the energy transfer efficiency between trophic levels, and reduce or slow down the carbon transfer to higher trophic levels in marine ecosystems. Recently, a study based on *in situ* experiments in the open ocean found that a range of predatory protists did not consume *Prochlorococcus*^20^. Therefore, it is rational to hypothesize that marine cyanobacteria (e.g. *Synechococcus* and *Prochlorococcus*) may employ evolutionary strategies to defend the predation of protists under the continuous predation pressure, which together with underlying mechanism(s) await experimental investigation.

In this study, to explore the anti-predation interactions between cyanobacteria and protists in the ocean, grazing experiments were conducted with representative marine picocyanobacteria and predatory protists, which lead to the observation of anti-predator defense of *Prochlorococcus* against *Uronema marinum*. Further mechanistic characterization showed that the *Prochlorococcus* specifically defend *U. marinum*’s predation through light-depend and inducible synthesis of anti-predator aldehydes. This anti-predation interaction between *Prochlorococcus* and *U. marinum* was shown to regulate the marine cyanobacterial growth and the *U. marinum* infection of marine fishes. To the best of our knowledge, this study provides the first report on the active anti-predation interaction between marine picocyanobacteria and protists, which together with the underlying mechanisms improve our understanding of the global element cycles and the stability of marine food chains and ecosystems.

## Results

### Grazing experiments reveal anti-predator defense traits of *Prochlorococcus* against *Uronema marinum*

To screen anti-predator defense traits from varied prey-predator interactions between cyanobacteria and protists in the ocean, grazing experiments were conducted with 4 representative marine picocyanobacterial strains (*Synechococcus* WH7803 and WH8102, and *Prochlorococcus* MED4 and NATL2A) and 3 predatory protists (*Uronema marinum, Pseudocohnilembus persalinus*, and *Cafeteria roenbergensis*) (Figure S1). The two protists (i.e. *P. persalinus* and *C. roenbergensis*) were shown to effectively prey on nearly all of the 4 picocyanobacterial strains, of which population dynamics aligned with the phase lag predicted by the classical prey-predator oscillation model ^21^. Specifically, the predatory protists’ cell concentrations increased by 6.8-354.6 times (mean, 116.8), and the 4 picocyanobacterial populations decreased by 43.3-100.0% (mean, 85.9%) accordingly (Figure S1A-D and F-H). One exception of the prey-predator interactions between the 2 protists and 4 cyanobacteria was that *Synechococcus* WH7803 evaded predation of *C. roenbergensis* through passive resistance, with only 3.3 ± 1.8% cell decrease of *Synechococcus* WH7803 and 5.6 ± 2.3% cell increase of *C. roenbergensis* (Figure S1E). Interestingly, different prey-predator interaction patterns were observed between the protist *U. marinum* and the two picocyanobacterial genera (*Synechococcus* and *Prochlorococcus*). For example, *U. marinum* effectively preyed on *Synechococcus* (WH7803 and WH8102), with 48.9-98.2% (mean, 74.0%) cell decrease of *Synechococcus* and 4.9-33.6 times (mean, 19.3) cell increase of *U. marinum* (Figure S1 I-J). In contrast, comparably fast growth of *Prochlorococcus* strains (MED4 and NATL2A) were observed in both grazing experimental sets with and without protist *U. marinum* (Figure S1K-L), indicating the anti-predator defense of *Prochlorococcus* against *U. marinum*. This anti-predator defense allowed the prey population to grow and persist at high cell density under predation pressure, being consistent with characteristics of a cost-free and active defense^22^.

To elucidate the anti-predator defense mechanism underlying prey-predator interactions between *Prochlorococcus* and *U. marinum*, both the *Prochlorococcus* MED4 and *U. marinum* were selected for subsequent grazing experiments. Time-series grazing experiments showed that *Prochlorococcus* MED4 grew to 1.5 × 10^8^ cells mL^−1^, being comparable to the MED4 cell concentrations in predator-free controls (Figure 1A). Accordingly, cell concentrations of *U. marinum* decreased by 92.8 ± 6.3% and remained unrecovered in subsequent 5 days (Figure 1B). These results suggested that *Prochlorococcus* MED4 employed an active defense mechanism to prevent protist’s predation by killing *U. marinum*. To further visualize the anti-predator defense process, both light and scanning electron microscopes were used to observe the details (Figure 1C-D; Movie S1; Figure S2A and B): stage-I, *U. marinum* ingested MED4 in food vacuoles via phagocytosis; stage-II, the contractile vacuoles of *U. marinum* cells started to swell after 30 minutes of MED4 ingestion; stage-III, the fast swelling of contractile vacuoles resulted in cytoplasmic content leakage within 18 seconds. In this process, the MED4-ingestion-triggered *U. marinum* cell lysis was accompanied with significant increase of toxic reactive oxygen species (ROS), compared to the experimental sets of ‘*U. marinum*’ and ‘*U. marinum* + *Synechococcus* WH8102’ (*p* < 0.05; Figure 1E; Figure S3). Notably, MED4 cell sizes changed with growth phases, i.e., 426 ± 69, 809 ± 193 and 685 ± 150 nm (in diameter) in lag-, exponential- and decline-phases, respectively (Figure S2C–E). To compare the MED4’s anti-predator defense activity in the three growth phases, protist

**Figure 1.**
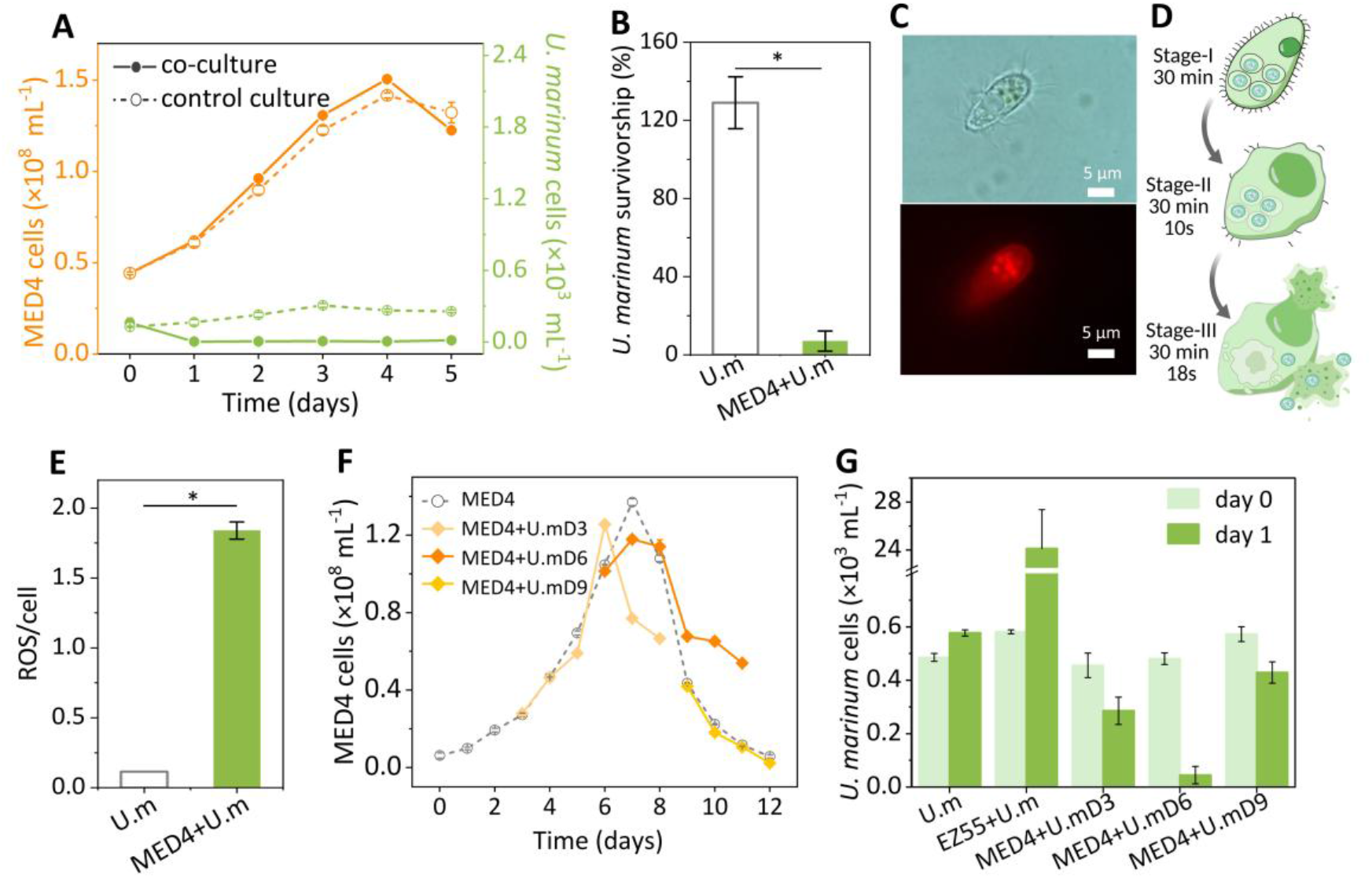
Predation defense of *Prochlorococcus* MED4 against *Uronema marinum*. (A) Temporal changes in cell density of MED4 (prey) and/or *U. marinum* (U.m; predator) in the MED4+U.m co-culture and in the MED4 and U.m control cultures. (B) The survivorship of *U. marinum* in the MED4+U.m co-culture and the U.m control culture after 24 hours incubation. (C) Light microscope images of *U. marinum* with ingested MED4 cells. Ingested MED4 cells are identified via chlorophyll autofluorescence (red). (D) Schematic diagram describing the three stages of MED4’s defense against *U. marinum* predation. (E) *U. marinum*-secreted ROS in the U.m culture and the MED4+U.m co-culture; ROS was measured in fluorescence and normalized to the *U. marinum* cell numbers. (F) Temporal changes in cell density of MED4 in cultures without *U. marinum* inoculation (MED4) and with *U. marinum* inoculation on day-3 (MED4+U.mD3), day-6 (MED4+U.mD6) or day-9 (MED4+U.mD9). (G) Cell density of *U. marinum* after its 0 and 1 day inoculation in the MED4+U.mD3, MED4+U.mD6 and MED4+U.mD9 experimental sets, as well as in the U.m control cultures with or without EZ55 amendment. Error bars represent SDs of triplicate experiments. Statistical significance is based on the T test (n = 3), which is denoted by asterisks (*), *p* < 0.05.

*U. marinum* was inoculated into three batches of MED4 cultures under the three different growth phases (Figure 1F). Specifically, in contrast to 41.4 ± 5.1 times of *U. marinum* cell increase through feeding *Alteromonas macleodii* EZ55, 40.7 ± 3.8%, 86.4 ± 6.7% and 25.1 ± 6.6% *U. marinum* cells were killed by MED4’s lag-, exponential- and decline-phase cells, respectively (Figure 1G), suggesting the highest anti-predator defense activity of exponential-phase cells of MED4.

### Light-dependent synthesis of anti-predator chemical(s) by *Prochlorococcus* to defend *Uronema*’s predation

For the active defense, preys generally use mechanical, chemical and/or biological weapons (e.g. T4SS/T6SS and virus) to harm and kill their predators^18^. In the anti-predation defense interactions between the *Prochlorococcus* and *Uronema*, both mechanical and biological weapons were excluded based on the fact that homologous genes were absent in *Prochlorococcus* genomes^23^ for encoding known mechanical secretion systems and lysogenic phages (Figure S4A; Table S1). Consequently, we hypothesized that MED4 employed the chemical weapon to defend *U. marinum* predation. To test this hypothesis, three sets of grazing experiments were prepared (Figure 2A-B): Set-I (MED4-filtrate), the *U. marinum* culture was fed with a bacterium (*Alteromonas macleodii* EZ55) and MED4 filtrate to test whether the culture filtrate of exponential-phase MED4 contained extra-cellular chemical(s) to defend *Uronema*’s predation; Set-II (freezing-MED4), the *U. marinum* culture was fed with freezing-killed MED4 cells to test whether *U. marinum* predated the freezing-killed exponential-phase MED4; Set-III (MED4+U.m-filtrate), the

**Figure 2.**
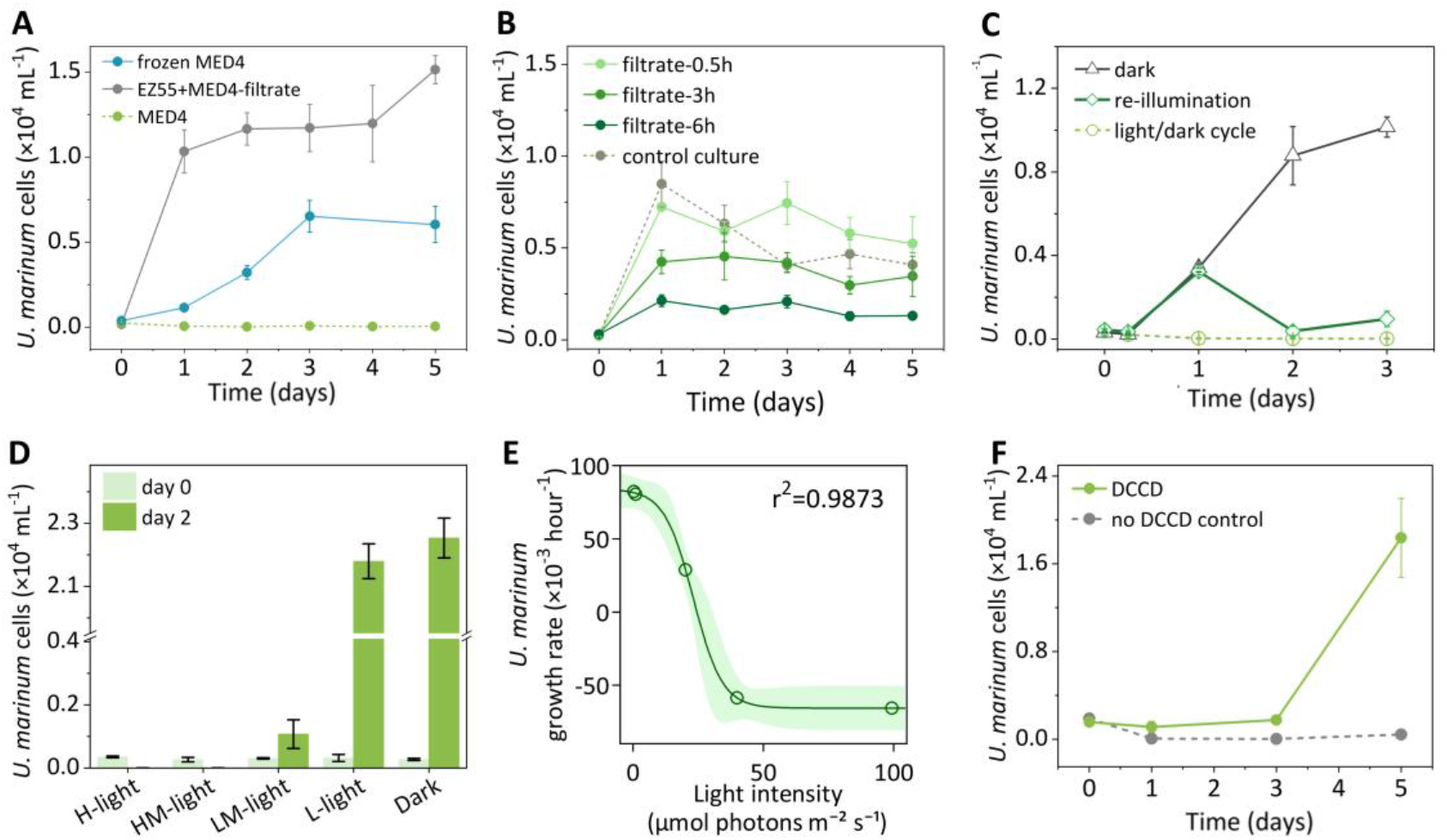
Inducible and light-dependent synthesis of anti-predator chemical(s) by *Prochlorococcus* to defend *Uronema*’s predation. (A) Temporal changes in cell density of *U. marinum* in U.m cultures amended with live MED4 cells (MED4), with freezing-killed MED4 cells (frozen MED4), or with live EZ55 cells and MED4 culture filtrate (EZ55+MED4-filtrate). (B) Temporal changes in cell density of *U. marinum* in EZ55+U.m co-cultures amended without (control culture) or with the MED4+U.m co-culture filtrate of 0.5 (filtrate-0.5h), 3 (filtrate-3h) and 6 hours (filtrate-6h) of co-cultivation. (C) Temporal changes in cell density of *U. marinum* in the MED4+U.m co-culture under light/dark cycle, continuous dark and re-illumination (2 days dark and then returned to the 2 days light/dark cycle) conditions. (D) Cell density of *U. marinum* after its 0 and 2 days’ inoculation in the MED4+U.m co-cultures under 5 different light intensities, i.e., 100 (H-light), 40 (HM-light), 20 (LM-light), 0.6 (L-light) and 0 (Dark) μmol photons m^−2^ s^−1^. (E) Correlation between the *U. marinum* growth rates and the light intensity based on the data shown in figure (D). Outer lines indicate the 95% confidence interval (*p*= 8.79 × 10−^10^). (F) Temporal changes of *U. marinum* cell density in MED4+U.m co-cultures with and without the ATP-synthesizing inhibitor DCCD. Error bars represent SDs of triplicate experiments.

*U. marinum* culture was fed with bacterium EZ55 and filtrate of MED4+U.m co-culture to test whether the chemical(s) for MED4’s defense was required to be induced by *U. marinum*. The high cell growth of *U. marinum* in Set-I and -II grazing experiments (i.e. 1.5 ± 0.8 × 104 and 6.5 ± 0.9 × 10^3^ cells mL^−1^ in Set-I and -II, respectively) suggested that exponential-phase MED4 cells without predation pressure did not synthesize intra- and extra-cellular chemicals to defend *U. marinum* predation (Figure 2A). In contrast, cell growth of *U. marinum* was obviously inhibited by amendment of *U. marinum*-exposed MED4 filtrate in the Set-III grazing experiments, and the peak cell concentrations of *U. marinum* decreased by 14.6 ± 0.8%, 50.0 ± 7.6% and 74.8 ± 3.7% with the *U. marinum*-exposed MED4 filtrate of 0.5, 3 and 6 hours’ *U. marinum* exposure, respectively (Figure 2B; Figure S4B). These results hinted that MED4 synthesized predation-induced chemical(s) to resist *U. marinum* predation. This inducible defense mechanism could be an adaptive strategy of the prey to be only activated upon sensing predation to balance the energy expenditure of survival^24^.

To confirm the energy-dependent inducible defense mechanism, the anti-predator activity of MED4 was tested with gradient light intensities and N,N-Dicyclohexylcarbodiimide (DCCD) as an ATP-synthesis inhibitor, based on the fact that synthesis of the inducible chemical(s) needed to be coupled with the energy metabolism of the photosynthetic MED4. Under a standard daily light-dark cycle, cell densities of *U. marinum* decreased from an initial number of 386 ± 28 cells mL^−1^ to 25 ± 7 cells mL^−1^ within 48 hours in the *Prochlorococcus*-*Uronema* grazing experiments. In contrast, under the continuous dark condition, *U. marinum* grew to 8.8 ± 1.4 × 10^3^ cells mL^−1^ by preying upon MED4 (Figure 2C). Notably, 87.9 ± 7.3% *U. marinum* cells were killed by MED4 after changing the dark condition to the standard daily light/dark cycles for 24 hours (Figure S4C). Further experiments on the correlation between light intensity and MED4’s anti-predator activity showed that the anti-predator activity was negatively correlated with light intensity, i.e., the growth rate of *U. marinum* decreased from 93.0 ± 3.1 × 10^−3^ to −92.1 ± 10.5 × 10^−3^ hour^−1^ when increasing light intensity from 0 to 100 μmol photons m^−2^ s^−1^ (r^2^ = 0.9873, Figure 2E). Similar with grazing experiments under the dark condition, DCCD-derived inhibition on ATP synthesis of MED4 cells significantly compromised the anti-predator activity of MED4 (*p* < 0.05), and *U. marinum* cells increased from 1.6 ± 0.1 × 10^3^ cells mL^−1^ in control experiments to 1.8 ± 0.4 × 104 cells mL^−1^ in DCCD-amended experiments (Figure 2F; Figure S4D-E). These results suggested that MED4’s synthesis of the anti-predation chemical(s) was energy-dependent and predator-inducible.

### *Prochlorococcus* employs aldehydes to defend *U. marinum* predation

To identify the inducible anti-predator chemical(s) of *Prochlorococcus*, transcriptomic and metabolomic analyses were performed with the *Prochlorococcus* MED4 culture and the MED4+U.m co-culture. In the transcriptomic analysis, the comparatively up*-*regulated genes in the MED4+U.m co-culture were mostly involved in energy metabolism, carbohydrate metabolism and metabolism of cofactors and vitamins, including the *mdh* and *queD* genes encoding methanol-to-formaldehyde and 7,8-dihydroneopterin 3’-triphosphate-to-acetaldehyde enzymes, i.e. methanol dehydrogenase (Mdh), 6-carboxy-5,6,7,8-tetrahydropterin synthase (QueD) (Figure S5A; Table S2). Further qPCR tests showed that the highest transcription of *mdh* and *queD* genes in the MED4+U.m co-culture significantly increased by 5.2 ± 0.6 and 1.9 ± 0.1 folds relative to them in the MED4 culture (Figure 3A; Figure S5B-C). Notably, transcription of the *mdh* and *queD* genes under the dark cultivation conditions of both MED4 and MED4+U.m cultures were comparable with, or even lower than, these in the normal MED4 culture, e.g. transcription of the *mdh* gene decreased by 19.0 ± 5.1% and 26.0 ± 9.1% in the MED4 and MED4+U.m cultures, respectively, under the dark condition, compared to that in normal MED4 culture (Figure 3A). Accordingly, GC-MS quantification showed 3.4- and 14.0-folds increase of formaldehyde and acetaldehyde concentrations, respectively, in the MED4+U.m co-culture relative to the MED4 culture (Figure 3B and Figure S5D). In contrast, only 1.4- and 4.7-folds increase in formaldehyde and acetaldehyde concentrations, respectively, were observed in the *P. persalinus*-exposed MED4 culture (MED4+P.p co-culture), compared to the MED4 cultures (Figure 3C). Therefore, the specifically promoted generation of formaldehyde and acetaldehyde in the MED4+U.m co-culture suggested their potential roles in MED4’s defense to *U. marinum* predation.

**Figure 3.**
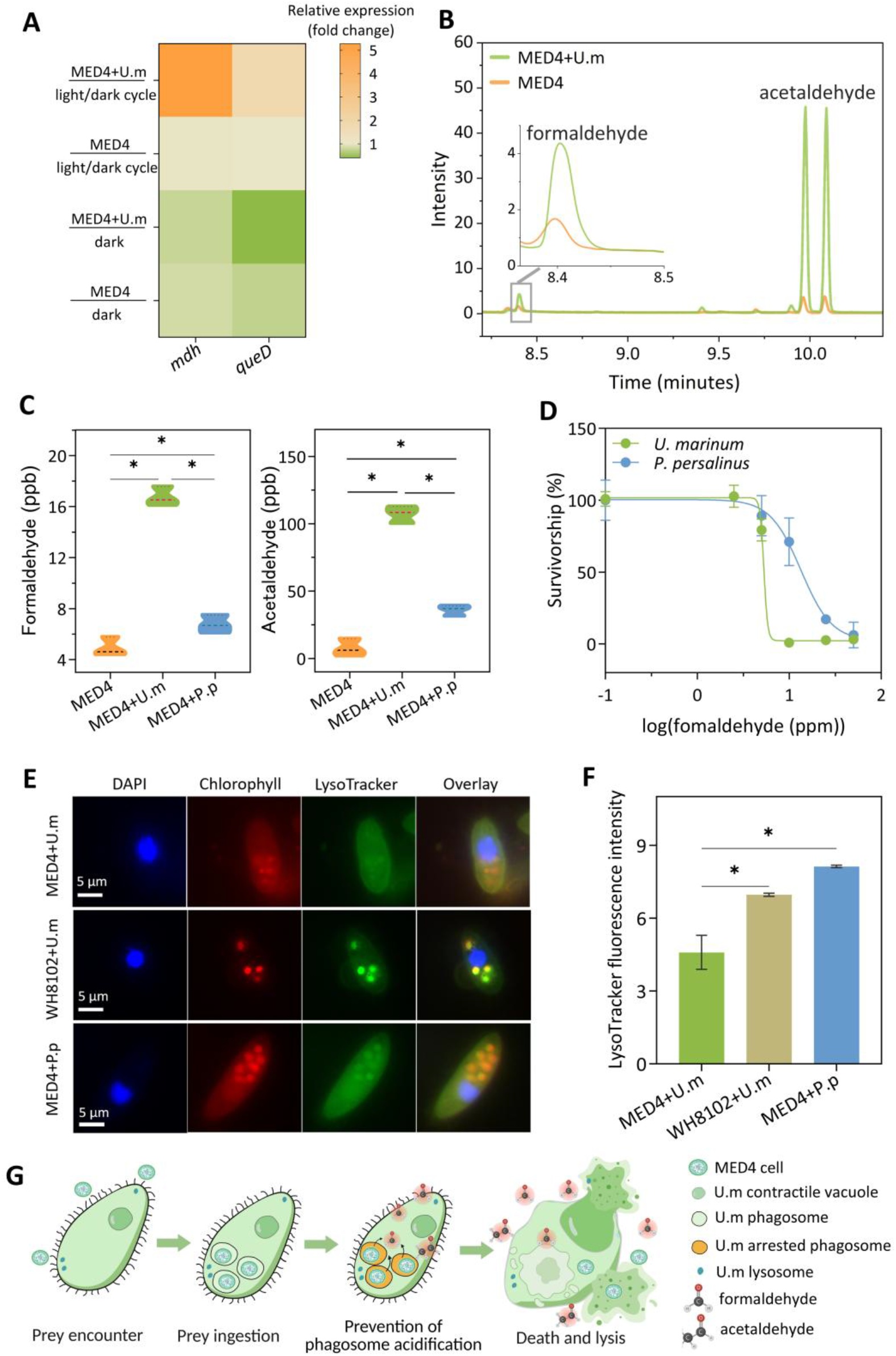
*Prochlorococcus* employs aldehydes to defend *U. marinum* predation. (A) Expression levels of Mdh (methanol dehydrogenase)- and QueD (6-carboxy-5,6,7,8-tetrahydropterin synthase)-encoding genes in the MED4+U.m co-cultures or MED4 cultures under light/dark cycle and dark conditions. (B) Overlaid GC-MS traces of aldehydes in the MED4 culture (orange) and the MED4+U.m co-culture (green). Retention times of formaldehyde and acetaldehyde were 8.4 and 9.9/10.1 minutes, respectively. (C) Peak concentrations of formaldehyde and acetaldehyde in the MED4 culture and in the MED4+U.m and MED4+P.p co-cultures. (D) Dose-response curves of formaldehyde against *U. marinum* and *P. persalinus*. (E) Fluorescence microscopy images of prey (MED4 or WH8102; red), predator cell nuclei (DAPI; blue), and acidification of predator phagolysosomes (LysoTracker; green) during predation by *U. marinum* (U.m) or *P. persalinus* (P.p). (F) LysoTracker fluorescence intensities to quantify phagolysosomal acidification in *U. marinum* or *P. persalinus* cells of the MED4+U.m, WH8102+U.m and MED4+P.p co-cultures. (G) Schematic diagram of aldehydes inducement in MED4 to defend *U. marinum* predation. Error bars represent SDs of triplicate experiments. Statistical significance is based on the one-way ANOVA followed by Tukey’s HSD post hoc test, which is denoted by asterisks (*), *p* < 0.05.

To further confirm aldehydes’ anti-predator activities, toxicity assessment was performed with the two groups of protists, i.e. *U. marinum* and *P. persalinus*. For the *U. marinum*, the 6-hour LC_50_ (median lethal concentration) of formaldehyde and acetaldehyde were 5.4 and 95.3 ppm, respectively (Figure 3D; Figure S5E), indicating the much higher toxicity of formaldehyde to *U. marinum* relative to acetaldehyde. In contrast, for the *P. persalinus*, the 6-hour LC_50_ of formaldehyde and acetaldehyde were 13.4 ppm and 244.5 ppm, respectively (Figure 3D; Figure S5F). In addition, the efficiency of digesting MED4 cells by the *U. marinum* and *P. persalinus* was compared by the tracking phagolysosome acidification with the LysoTracker Red DND-26 that specifically stained prey-susceptible acidic organelles in live cells (Figure 3E-F, Figure S5G-H). Staining results showed the obvious inhibition of phagolysosomal acidification and digestion and consequent accelerated aldehyde accumulation in the *U. marinum*, compared to the effective digestion of MED4 by *P. persalinus* using the phagolysosomal acidification. These toxicity tests and phagolysosomal acidification results, together with observations of the much higher amounts of induced formaldehyde and acetaldehyde in *U. marinum*-relative to *P. persalinus*-exposed MED4 cultures, worked together to generate a scenario to reveal how *Prochlorococcus* employed aldehydes (particularly the formaldehyde) to specifically defend *U. marinum* predation (Figure 3G): first, the exposure of *Prochlorococcus* MED4 to *U. marinum* induced the generation of formaldehyde as a primary anti-predator chemical; then, the phagocytosis of MED4 cells by formaldehyde-sensitive *U. marinum* into food vacuoles, where MED4 cells were not efficiently digested, led to intracellular aldehyde accumulation and magnified aldehyde toxicity (relative to extracellular aldehydes in the medium), and resulted in lethality of *U. marinum*. In contrast, fewer aldehydes were generated in *P. persalinus*-exposed MED4, and the ingested MED4 cells were efficiently digested through lysosomal acidification within *P. persalinus* to minimize intracellular aldehyde accumulation, which together with comparatively higher tolerance of *P. persalinus* to aldehydes, rationalized the observation of no anti-predator activity of MED4 to *P. persalinus* predation.

### Impact and implications of the *Prochlorococcus*-*U.marinum* anti-predation interactions

The *Prochlorococcus*’s anti-predator activity could have a range of impact and implications, including its regulation of global *Prochlorococcus-Uronema* distribution and carbon/energy flow in the marine food web, as well as its control of *U. marinum*-derived parasitic diseases of marine fishes. Totally 772 metagenomes and 16S/18S rRNA gene amplicon sequencing datasets were collected to assess the global *Prochlorococcus-Uronema* distribution pattern (Figure 4A). Results showed the mutually exclusive distribution of *Prochlorococcus* and *Uronema* in marine environments, which was further confirmed with multiple exclusivity indices, i.e., 0.979 Jaccard distance and 0.959 Sørensen dissimilarity (Figure 4B). The *U. marinum* as a common and deadly parasite of marine fishes primarily infected fish skin, gill and muscle tissue, of which infection control in fish farms generally referred to pesticides, antibiotics and copper treatment, and consequently raised concerns in secondary aquatic contamination and food safety^25, 26^. To prevent these chemical treatment-derived side effects, cyanobacterium *Prochlorococcus* MED4 could be used as fish feed supplement to control the *U. marinum* infection of marine fishes (i.e. fish *Chromis viridis* in this study). Results showed that MED4 supplement for 4 days reduced *U. marinum* infection by 72.5 ± 16.7% and 83.8 ± 6.0% in the fish gill and muscle tissues, respectively, relative to the experimental set of *U. marinum-*infected fishes without supplementing MED4 (Figure 4C). These results indicated the promising application of *Prochlorococcus* as a biocontrol agent in treatment of *U. marinum* infection in marine fishes.

**Figure 4.**
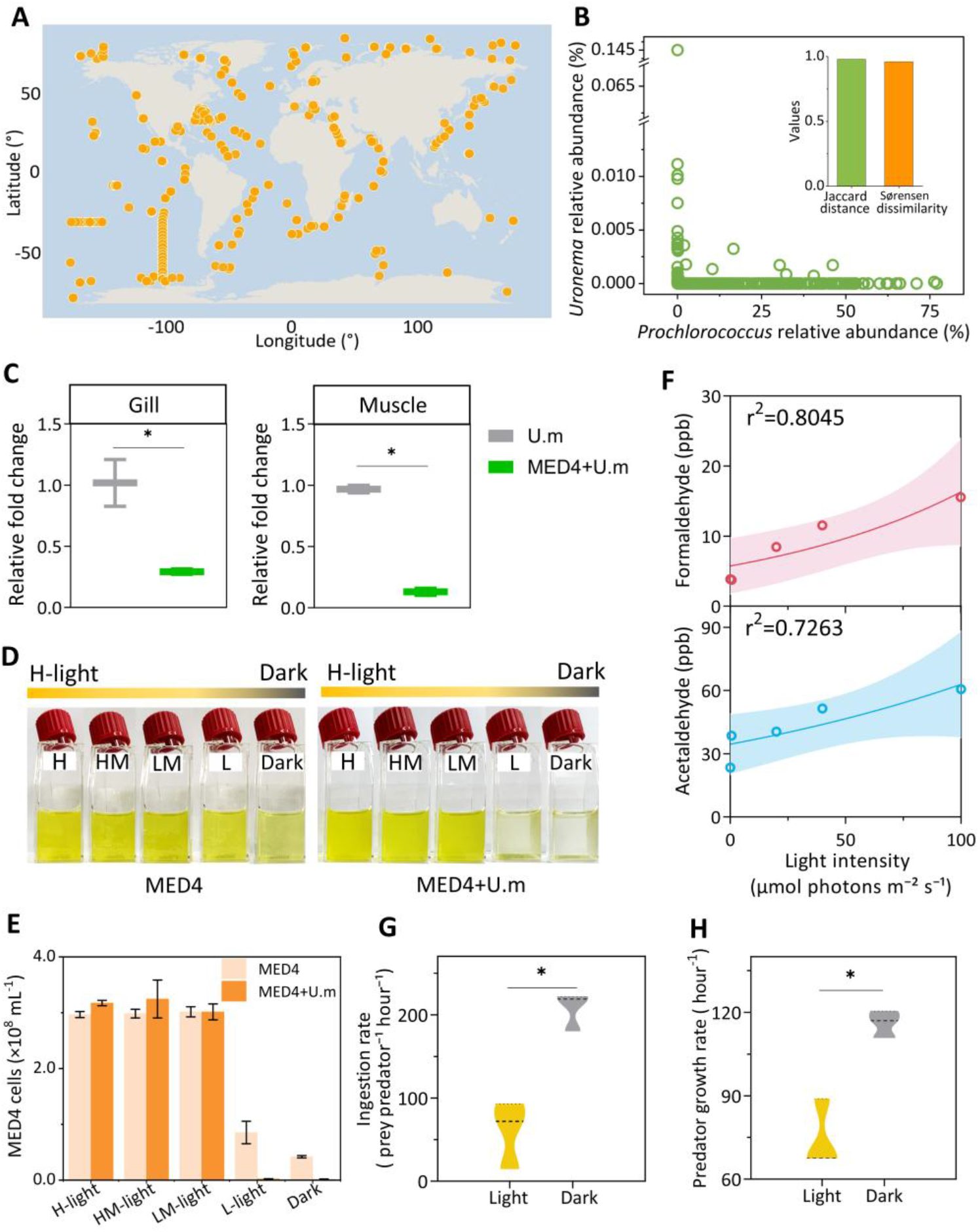
Implications of the *Prochlorococcus*’s anti-predation defense against *Uronema marinum*. (A) Global marine microbial community data (including metagenomic and 16S/18S rRNA gene amplicon sequencing data; n = 772) for analyzing the co-occurrence of *Prochlorococcus* and *Uronema*. (B) Relationship between *Prochlorococcus* and *Uronema* based on their relative abundance in the global marine microbial communities, and their mutually exclusive occurrence assessed with Jaccard distance and Sørensen dissimilarity indices. (C) Abundance changes of *U. marinum* in gill and muscle tissues of marine fish *Chromis viridis* with and without amendment of MED4. Cultivation of MED4 cultures and MED4+U.m co-cultures on day-2 (D) and associated MED4 cell density (E) under growth conditions with 5 gradient light intensities, i.e. 100 (H), 40 (HM), 20 (LM), 0.6 (L), and 0 (Dark) μmol photons m^−2^ s^−1^. (F) Correlation between light intensity and produced aldehyde concentration (formaldehyde, top panel, *p*=0.011; acetaldehyde, bottom panel, *p*=0.045). Outer lines indicate 95% confidence interval. Impact of light condition (day and night) on the ingestion rate (G) and growth rate (H) of *P. persalinus* in the MED4+P.p co-culture. Error bars represent SDs of triplicate experiments. Statistical significance is based on the T test (n = 3), which is denoted by asterisks (*), *p* < 0.05.

The light-dependent and inducible synthesis of aldehydes in *Prochlorococcus* cells upon *U. marinum* predation provided clues to understand and mediate interactions between cyanobacteria and protists. Decreasing light intensity was shown to promote protists’ predation on the cyanobacteria, which provided a strategy to improve the control of cyanobacterial growth. In this study, 99.3 ± 0.1% and 99.5 ± 0.1% less cells of *Prochlorococcus* MED4 (Figure 4 D-E), together with obvious lower aldehyde concentrations, were observed in the MED4+U.m co-culture under low-light (0.6 μmol photons m^−2^ s^−1^) and dark conditions, respectively, compared to its normal light condition (20-100 μmol photons m^−2^ s^−1^; Figure 4F and Figure S6). Moreover, interestingly, in the MED4+P.p co-cultures, lower ingestion and growth rates were observed under the normal light condition, relative to the dark condition (Figure 4G-H), which could be due to the comparatively lower amount of synthesized aldehydes of *Prochlorococcus* MED4 under the dark condition (Figure 4F). Therefore, improved predation efficiencies could be expected to control cyanobacterial growth by supplementing protists in night time (dark condition) or by adding cover-materials to overlay the water surface (low-light condition).

## Discussion

This study reports the first observation of, as well as mechanistic insights into, *Prochlorococcus* anti-predator activity to defend *U. marinum* predation, which may increase the *Prochlorococcus*’s fitness in natural selection and consequently have a global impact on the carbon/oxygen cycles based on the fact of *Prochlorococcus* as the most abundant photosynthetic and oxygenic microorganisms in the ocean. The success of *Prochlorococcus* anti-predator defense against *U. marinum* predation exclusively depends on three things, i.e. light intensity, predation pressure-induced aldehydes and low lethal concentrations of aldehydes to the predator (*U. marinum*). Previous studies have shown that exposure of cyanobacteria to grazing cues enhances the photosynthetic activity, including quantum yield of photosystem II (PSII) and the maximum electron transport rate^27^. Moreover, a diel rhythm was observed in picocyanobacteria’s grazing of heterotrophic microzooplankton, i.e. comparatively low and high grazing rates presented in the day and night time, respectively^28^. Similarly, harmonic oscillations in *Prochlorococcus* cell production and mortality rates were observed in the subtropical gyre, with a peak in mortality consistently occurring around 6 hours after dusk^29^. Nonetheless, reasons behind these observations were unknown. Our study rationalizes these observations by providing mechanistic insights into the light-dependent inducible synthesis of anti-predator aldehydes in *Prochlorococcus* cells upon protists’ predation.

The long-term prey-predator interactions in the marine food web and associated microbial loop may trigger ‘arms race’ among picophytoplankton, bacteria and microzooplankton, which further affect the carbon and energy flows and ecosystem stability through the marine food web^30^. For example, the distribution of *Prochlorococcus*, a major contributor in the microbial loop to the marine carbon and energy flows, was determined primarily by predatory relationships instead of previously assumed temperature^31^. In the traditional marine food web (black lines in Figure 5), carbon and energy linearly flow through phytoplankton, protozoans, zooplankton and carnivores (grazing food chain), in which virus and bacteria can recycle carbon/energy resources by lyzing bacterial/cyanobacterial cells and degrading the food-chain-derived organic carbon to form the viral loop (or viral shunt) and microbial loop, respectively^32, 33^. Recently, bacteria were identified to employ anti-predator defense strategies to recirculate the carbon/energy flows in the marine food web^34^, which complemented the traditional marine food web by channeling the carbon flow from protozoans/zooplankton back to heterotrophic microorganisms. Our identification of the anti-predator activity of *Prochlorococcus* against *U. marinum* predation add a new connection between the heterotrophic microorganisms and protozoans/zooplankton, highlighting the contribution of *Prochlorococcus*’s anti-predator activity to the carbon/energy backflow in the marine food web (red lines in Figure 5). The new comprehensive marine food web will advance our understanding of the role of marine primary production and improve predictions of marine carbon sequestration and ecosystem resilience under the global climate change.

**Figure 5.**
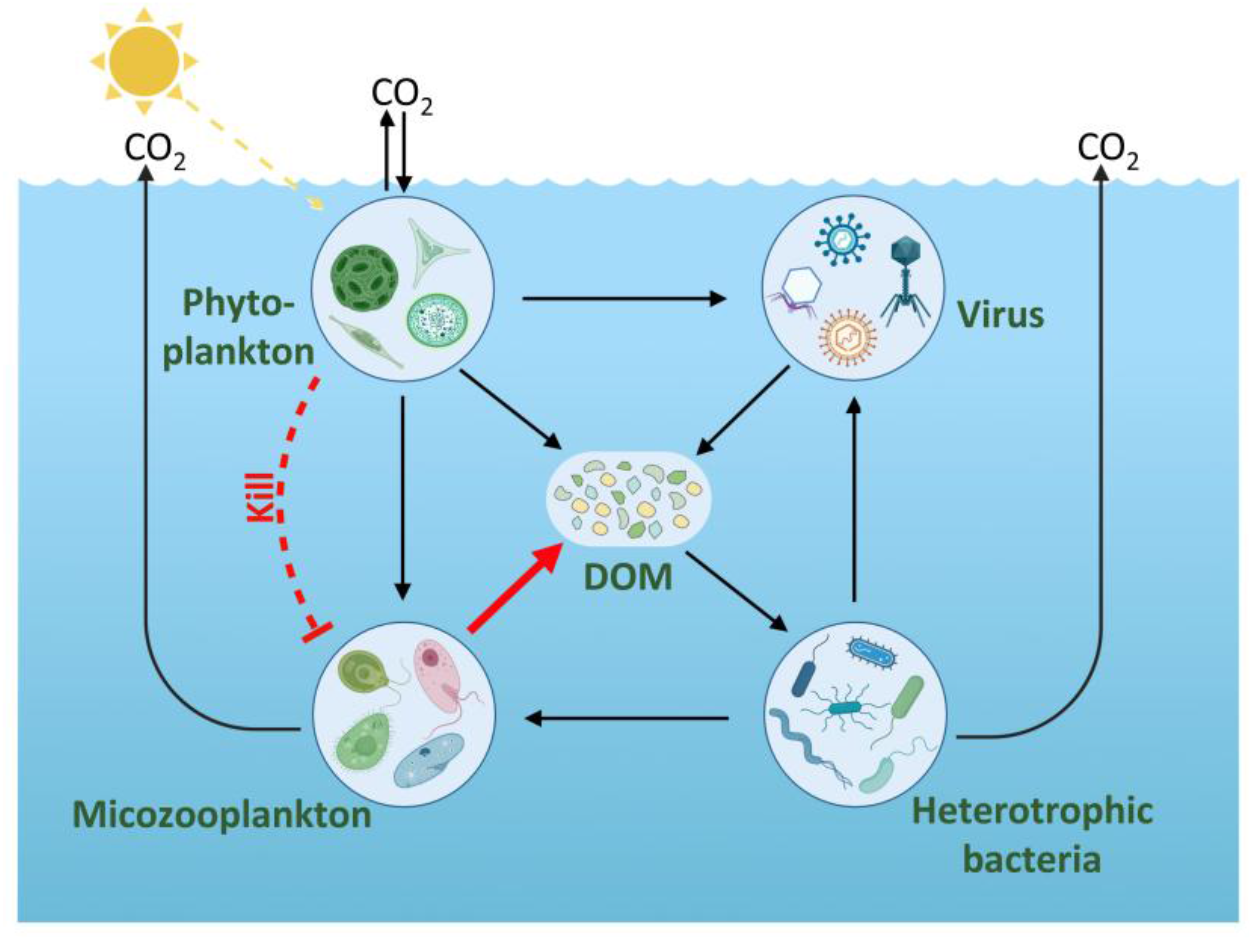
Microbial loop–mediated carbon fluxes modulated by the *Prochlorococcus*-*Uronema* (microzooplankton-phytoplankton) anti-predation interaction. Carbon transfer within the marine microbial loop occurs among phytoplankton, microzooplankton, viruses and heterotrophic bacteria through key interactions/processes including grazing, viral lysis and DOM recycling (black line). The anti-predation interaction between microzooplankton and phytoplankton identified in this study can modulate these carbon fluxes through the anti-predation derived DOM (dissolved organic matter) release from microzooplankton (red line).

## Acknowledgments

This study was financially supported by the Southern Marine Science and Engineering Guangdong Laboratory (Zhuhai) (SML2021SP317 to S.W. and SML2024SP022 to Z.H.) and National Natural Science Foundation of China (42161160306 to S.W. and 42430707 to Z.H.).

## Materials and methods

### Experimental model organisms

#### Picocyanobacterial strains

Marine cyanobacterial strains, including *Prochlorococcus* NATL2A^35^, *Prochlorococcus* MED4^36^, *Synechococcus* WH8102 and *Synechococcus* WH7803^37^, were cultured in AMP1 medium^38^ at 22 °C with a daily 14 h: 10 h light: dark cycle. *Prochlorococcus* NATL2A was a low-light adapted strain, and cultured at 20 μmol photons m^−2^s^−1^ The other three strains were high-light adapted and maintained at 70 μmol photons m^−2^ s^−1^. To ensure axenicity, cyanobacteria were routinely tested in following sterile broths, i.e. ProAC^39^, MPTB^40^, or ProMM^41^.

#### *Alteromonas macleodii* EZ55

*Alteromonas macleodii* EZ55, a marine heterotrophic bacterium, was cultured in the ProAC medium at 30 °C with 200 rpm shaking as described^39^, of which freezing-killed cells were provided as a food source (prey) to support rapid cell growth of protists (predator). The freezing-killed EZ55 cells were prepared through being frozen at −80 °C overnight and then thawed for 1 hour.

##### Protists strains Protists

Two marine heterotrophic ciliates (*Uronema marinum* and *Pseudocohnilembus persalinus*) and one heterotrophic nanoflagellate (*Cafeteria roenbergensis*) were maintained in autoclaved artificial seawater supplemented with sterile rice grains. These protists were isolated from coastal waters of the Hong Kong and Xiamen cities in China, which were widely distributed in subtropical regions^42,43^.

##### Chromis viridis

Marine fish, *Chromis viridis*, was purchased from a local fish market in Guangzhou (Guangdong, China) and maintained in a recirculating aquaculture system. The culture was supplied with fully aerated artificial seawater under a strictly controlled condition, i.e., 22.0 ± 1.0 °C water temperature, 33.0 ± 1.0‰ salinity, and a daily 14 h:10 h light-dark (diel) cycle. All experiments performed in this study adhere to the American Physiological Society’s Guiding Principles in the Care and Use of Animals and were approved by the Institutional Animal Care and Use Committee of Sun Yat-sen University (SYSU-IACUC-2026-B1259).

### Grazing experiments

Grazing experiments were conducted with the four picocyanobacterial strains (*Prochlorococcus* NATL2A and MED4, *Synechococcus* WH8102 and WH7803) as preys and three protists (*U. marinum* and *P. persalinus* and *C. roenbergensis*) as predators/grazers. Grazer cells were introduced into the prey cultures to reach a final prey:grazer ratio of 3×104 for the nanoflagellate *C. roenbergensis* and 3×10^5^ for the ciliates *U. marinum* and *P. persalinus* (0.1-2.0×10^8^ prey cells mL^−1^). Initial grazer concentration was 1-6 × 10^3^ cells mL^−1^ for the nanoflagellates and 1-6 × 10^2^ cells mL^−1^ for the ciliates, and the grazing assays were conducted for 48 hours. Samples were collected for cell counts at t=0 and 48 hours. Based on the observation of MED4’s anti-predation against *U. marinum* in initial-screening grazing assays, 5-day grazing assays (MED4, ~5 × 10^7^ cells mL^−1^; a final prey:grazer ratio of 3×10^5^ for the ciliate *U. marinum*) were conducted to monitor their population dynamics. Control cultures of grazers without prey or prey without grazers were included. Daily samples were collected for cell enumeration.

To investigate the effect of prey growth phase on grazing resistance in MED4+U.m co-cultures, the *U. marinum* cultures (1-6 × 10^2^ cells mL^−1^) were subjected to three dietary treatments, i.e. lag-(day 3; MED4+U.mD3), exponential-(day 6; MED4+U.mD6), and decline-phase (day 9; MED4+U.mD9) MED4 cells. MED4 control culture was prepared with MED4 inoculation only. Two U.m control cultures were prepared with (EZ55+U.m) and without EZ55 feeding (U.m). Samples were collected for cell counts at 24-hour intervals and the survivorship of *U. marinum* was quantified at t = 24 hours as described as (*N*_t_ / *N*_0_) ×100%, where *N*_t_ and *N*_0_ represent the grazer densities at 24 and 0 hours, respectively.

To test the effect of predator’s presence on the active defense of MED4, the *U. marinum* cultures (1-6 × 10_2_ cells mL^−1^) were subjected to four groups of dietary treatments: (1) MED4, exponential growth phase MED4 cells were provided as prey with a final cell density of 5 × 10^7^ cells mL^−1^; (2) frozen MED4, MED4 cells (nominal concentration, 1 × 10^8^ cells mL^−1^) were frozen at −80 °C overnight and then thawed for 1 hour to feed *U. marinum* cultures; (3) EZ55+MED4-filtrate, freezing-killed EZ55 cells were provided as prey (1 × 10^8^ cells mL^−1^, being frozen at −80 °C overnight and then thawed for 1 hour) in exponential-phase MED4 culture filtrate; (4) EZ55+(MED4+U.m-filtrate), freezing-killed EZ55 cells were provided as prey (1 × 10^8^ cells mL^−1^, being frozen at −80 °C overnight and then thawed for 1 hour) in filtrate of the MED4+U.m co-culture with 0.5 (filtrate-0.5h), 3 (filtrate-3h) and 6 (filtrate-6h) hours co-cultivation. The U.m control culture was prepared with *U. marinum* and no prey feeding.

To compare the grazing efficiency in MED4+P.p co-cultures under light or dark conditions, the *P. persalinus* cultures (1-6 × 10^2^ cells mL^−1^) were subjected to two cultivation conditions, i.e., light (mimicking day condition; 70 μmol photons m^−2^ s^−1^) and dark (mimicking night condition; 0 μmol photons m^−2^ s^−1^). MED4 control culture was prepared with MED4 inoculation only. In the MED4+P.p co-cultures and MED4 control cultures, MED4 cells were amended with a final concentration of ~5×10^7^ cells mL^−1^. The *P. persalinus* control cultures were prepared with *P. persalinus* inoculation only. Samples for cell count were collected at 0 and 10 hours after the co-culture establishment. The grazer’s growth rate (μ, hour^−1^) and ingestion rate (prey cell^−1^ hour^−1^; based on the change in prey density over time and normalized predator cells) were calculated for the grazing experiments under the ‘light’ and ‘dark’ conditions as described^44^.

### Impact of light intensity and ATP-inhibition on grazing resistance

To quantify the effect of light conditions on grazer cell growth, U.m cultures (1-6 × 10^2^ cells mL^−1^) were divided into three distinct illumination treatments: (1) dark, continuous darkness for 4 days; (2) re-illumination, complete darkness for 2 days followed by a standard light/dark (14h/10h) cycle condition for 2 days; (3) light/dark cycle, a standard light/dark cycle for 4 days. Furthermore, to test the effect of light intensities on grazer cell growth and predation resistance, the U.m cultures were divided into five distinct illumination treatments for 2 days, i.e., high light (H-light, 100 μmol photons m^−2^ s^−1^), moderately high light (HM-light, 40 μmol photons m^−2^ s^−1^), moderately low light (LM-light, 20 μmol photons m^−2^ s^−1^), low light (L-light, 0.6 μmol photons m^−2^ s^−1^), and complete darkness (Dark, 0 μmol photons m^−2^ s^−1^). The exponential-phase MED4 cells were introduced at a final density of ~5×10^7^ cells mL^−1^ in each treatment. For all illumination experiments, all treatments were prepared in triplicates with T25 flasks. Daily samples for grazer cell count were fixed with 10% Lugol’s solution in a 96-well plate. *U. marinum* cells were counted by using an inverted microscope. To test the effect of ATP inhibition on MED4’s anti-predation activity, the ATP synthase inhibitor *N,N*’-dicyclohexylcarbodiimide (DCCD) was added to the MED4+U.m co-culture at a final concentration of 0.01 μM, of which the concentration was confirmed to be non-cytotoxic to MED4 and *U. marinum* cells. The specific growth rate (μ, hour^−1^) of *U. marinum* was calculated for both the control and DCCD treatments. Additionally, MED4 cell viability was assessed using DMAO and Hoechst33342, of which staining images were captured using a fluorescence microscope (ECHO; San Diego, CA, USA).

### MED4 treatment of *U. marinum*-infected *Chromis viridis*

To evaluate the efficacy of MED4 treatment of *U. marinum*-infected marine fish, an experimental infection was established using *Chromis viridis*, based on its susceptibility to *U. marinum*^25, 26^. Healthy *C. viridis* individuals were selected with an average initial body weight of 1.3 ± 0.1 g and an average body length of 4.5 ± 0.2 cm. All fish were acclimated under the laboratory condition for one week before being used in the formal experiments. Following the acclimation, the experimental infection trial was conducted by exposing healthy *C. viridis* to *U. marinum* (100 cells mL^−1^) for 5 days. On day 5 prior to the treatment, three randomly selected fish were humanely euthanized using MS-222 (100 mg L^−1^) to verify the successful establishment of the infection. Microscopic examination of skin and gill scrapes from these individuals confirmed the presence of live *U. marinum*. Subsequently, the infected fish were randomly assigned to two experimental groups (n = 3 per group) to evaluate the therapeutic efficacy: (1) MED4+U.m, a 4-day treatment group receiving MED4 cells with a final concentration of 1.0 × 10^7^ cells mL^−1^; and (2) U.m, an untreated positive control group receiving no MED4 cells. A negative control group (n = 3) consisted of healthy and uninfected fish maintained under identical conditions but without exposure to *U. marinum* or MED4 cells. During the totally 9 days trial, approximately 20% of the seawater in all tanks was exchanged daily with artificial seawater and MED4 cells were proportionally replenished in the MED4+U.m group to sustain the target therapeutic concentration. Upon completion of the experiment, fishes were euthanized with an overdose of MS-222 (100 mg L^−1^). Gill and muscle tissues were harvested and immediately flash-frozen in liquid nitrogen before storage at −80 °C. Genomic DNA was extracted using the Marine Animal Tissue DNA Kit (Tiangen; Beijing, China). The relative parasite burden of *U. marinum* was determined by real-time quantitative PCR (qPCR) targeting the cytochrome-c oxidase subunit 1 (*cox1*) gene. Data were analyzed using the ΔΔCt method, with the *18S rRNA* gene serving as the endogenous control for normalization.

### Intracellular ROS measurements

The dye 2′,7′-Dichlorofluorescin diacetate (DCF-DA, Solarbio; Beijing, China) was used to quantify intracellular reactive oxygen species (ROS) in *U. marinum*. The *U. marinum* cells from U.m cultures, MED4+U.m and WH8102+U.m co*-*cultures were harvested by filtration and washed three times with sterile seawater. Cells were resuspended in fresh seawater containing 5 μM DCF-DA and incubated in the dark for 30 minutes. Fluorescence emission at 525 nm (excitation at 488 nm) was recorded using a microplate reader (Varioskan LUX, Thermo Fisher; Waltham, MA, USA). Measurements across plates were normalized by calculating the dispersion of empty and blank measurements per plate from the overall mean and subtracting the difference from the respective fluorescence measurements and then normalized to the cell numbers per milliliter^45^.

### Antiprotozoal activity testing and LC50 determination

*U. marinum* and *P. persalinus* were used to assay antiprotozoal activity of formaldehyde and acetaldehyde. *U. marinum* and *P. persalinus* cultures were adjusted to an initial density of 500 cells/mL in 10 mL of artificial seawater and exposed to gradient concentrations of formaldehyde or acetaldehyde. *U. marinum* and *P. persalinus* cultures without the addition of formaldehyde or acetaldehyde were prepared as negative controls. Following a 6 h incubation at 22°C, samples were fixed with Lugol’s solution at a final concentration of 10% for microscopic enumeration. Ciliate densities were measured and cell survivorship was calculated as a percentage relative to the negative controls. The survivorship values were plotted against the decadic logarithm of the aldehyde concentration and the LC50 value was determined using PRISM (GraphPad, v10).

### Light and fluorescence microscopy analyses

To monitor predation dynamics, live cells from the MED4+U.m co-culture and the U.m control culture were examined. The initial density of MED4 was adjusted to ~5 × 10^7^ cells mL^−1^, while *U. marinum* was inoculated at a density of ~500 cells mL^−1^. Movies and real-time observations were acquired using the light microscope (ECHO; San Diego, CA, USA) under 200× magnification.

For the fluorescence microscopy analysis, *U. marinum* (~500 cells mL^−1^) was co-incubated with MED4 (final density, ~5 × 10^7^ cells mL^−1^) for 0.5 hours and subsequently fixed with 2.5% (v/v) glutaraldehyde. To confirm predation, ingested MED4 cells were subsequently identified within the *U. marinum* cell food vacuoles by overlaying brightfield images with MED4’s red autofluorescence.

To assess the intracellular digestion process, two ciliate cultures (~500 cells mL^−1^) were subjected to three treatments with prey cells (~5 × 10^7^ cells mL^−1^): (1) MED4+U.m, *U. marinum* cells were provided with MED4 as prey; (2) WH8102+U.m, *U. marinum* cells were fed with WH8102; (3) MED4+P.p, *P. persalinus* cells were provided with MED4 as prey. The control cultures were prepared with *U. marinum* or *P. persalinus* cultures without prey feeding. Once the prey-ciliate interaction was initiated, subsamples of the ciliates after 0.5 hours of contact time were collected and fixed with 2.5% (v/v) glutaraldehyde and then stained with DAPI (5 μg mL^−1^) to visualize the ciliate nucleus. Ciliate cells were treated with LysoTracker Green DND-26 (75 nM final concentration) for 30 minutes to stain the acidic lysosomal compartments to observe the digestion process using fluorescence microscopy (ECHO; San Diego, CA, USA). Lysosomal digestion intensity was quantified using ImageJ software (NIH; Bethesda, MD, USA).

### Scanning electron microscopy

To characterize the cell morphology and size of MED4 at different growth phases, cell images were captured by scanning electron microscopy (SEM, Sigma 500; Zeiss, Oberkochen, Germany). Additionally, SEM was utilized to observe the interaction between *U. marinum* and MED4. Culture samples for the SEM analysis were fixed with 2.5% (w/v) glutaraldehyde and stored at 4°C overnight. Then, the samples were filtered through 0.2 μm isopore Millipore membranes, dehydrated using an ethanol gradient series (30%, 50%, 70%, 85%, 90%, and 100%) before being cleaned with butanol and dried. Images were collected in backscattered and secondary electron modes using a Zeiss Sigma 500 FEG-SEM operating at 3 kV^46^.

### Enumeration of prey and predators

Prey cells were quantified with flow cytometry (Accuri C5, BD Biosciences; San Jose, CA, USA) using glutaraldehyde-fixed samples (2.5% final concentration). Cyanobacterial populations were identified by plotting chlorophyll fluorescence (excitation 488 nm, emission 680 nm) against side scatter. Grazer cells were fixed with 10% Lugol’s solution in a 96-well plate and counted under an inverted microscope (ECHO; San Diego, CA, USA).

### GC-MS analysis of aldehydes

*Prochlorococcus* MED4-derived aldehydes (formaldehyde and acetaldehyde) were measured using headspace solid-phase microextraction coupled with gas chromatography/mass spectrometry (HS-SPME-GC/MS) as described^47, 48^. Specifically, 30 mL of culture samples were placed in a sealed vial, spiked with the internal standard 1,2-dibromopropane (50 mg L^−1^) and PFBHA derivatization reagent (50 mg mL^−1^), vortex-mixed, and incubated at 60 °C for 1 hours. After cooling to the room temperature, aldehydes were extracted by exposing a pre-conditioned (260 °C, 60 minutes) CAR/PDMS/DVB-coated SPME fiber to the vial headspace at 80 °C for 30 minutes. The fiber was then thermally desorbed in the GC injector. Chromatographic separation was performed with a Rtx-5MS column (30 m × 0.25 mm, 0.25 μm film; Restek, Bellefonte, PA, USA) with helium as a carrier gas at a constant flow of 1.2 mL minutes^−1^. The oven temperature program was: 40 °C (hold 2 minutes), ramped at 10 °C/ minutes to 180 °C, then at 20 °C/ minutes to 240 °C (hold 3 minutes). Mass spectrometric detection was conducted in electron-ionization (EI) mode at 70 eV, and target analytes were quantified using selected-ion-monitoring (SIM) mode.

### Genome annotation and comparative analyses

The genomes of *Prochlorococcus* MED4 (GenBank accession number, GCA_000011465.1), *Prochlorococcus* NATL2A (GCA_000012465.1), *Synechococcus* WH7803 (GCA_000063505.1), and *Synechococcus* WH8102 (GCA_000195975.1) were collected from the National Center for Biotechnology Information (NCBI) database. Functional annotation of these genomes was carried out with eggNOG-mapper v2.1.9^49^. To compute the pangenome, an ‘anvi’o genomes storage’ database was first generated from the FASTA files of the four genomes using the ‘--internal-genomes’ flag. Subsequently, the program ‘anvi-pan-genome’ was run with the generated storage database, along with the flag ‘--use-ncbi-blast’ and the parameters ‘--minbit 0.5′ and ‘--mcl-inflation 10′. Hierarchical clustering analyses of gene clusters (based on their distribution across genomes) and genomes (based on shared gene clusters) were performed using Euclidean distance and Ward clustering by default. The results were visualized using the ‘anvi-display-pan’ program^50^.

### Transcriptome sequencing and RT-qPCR

Transcriptomic analysis was performed for the MED4 cultures and MED4+U.m co-cultures as described ^51^. Briefly, samples for total RNA extraction were collected by centrifugation (15 minutes, 12,000× g, 4 °C) from mid-exponential phase cultures. RNA samples were then mixed with TRIzol reagent (Invitrogen). Total RNA was extracted with HiPure Universal RNA Mini Kit (Magen; Guangzhou, China), from which fragmented DNA was removed using DNase I. The RNA-Seq libraries were constructed from the rRNA-depleted RNA samples using the ALFA-SEQ RNA Library Prep Kit II (Vazyme; Nanjing, China) and then sequenced on an Illumina Novaseq 6000 platform. The RNA-Seq library construction and sequencing services were provided by Magen Biotechnology (Guangzhou, China). RNA-Seq raw data in fastq format were quality-filtered and trimmed using Trimmomatic (v.0.35)^52^ to acquire the clean reads. To eliminate ribosomal RNA contamination, clean reads were aligned to the NCBI Rfam database by Bowtie2 (v2.33)^53^ and aligned sequences were discarded. The remaining mRNA sequences were mapped to the reference genome of *Prochlorococcus* MED4 by Bowtie2. Gene expression levels were quantified as transcripts per million (TPM) described earlier^54^.

For gene expression analysis, MED4 cells were cultured with or without *U. marinum* under light and dark conditions for 0.5, 3 and 6 hours. RNA isolation was performed as described in the ALFA-SEQ Magnetic Universal RNA Kit (Findrop). The resulting RNA was reverse-transcribed into cDNA using ALFAFast King RT Mix with gDNA Remove (Findrop) under the following condition, i.e., 37°C for 2 minutes, 50°C for 10 minutes, and 80°C for 2 minutes. Quantitative RT-PCR was carried out with cDNA, gene-specific primers (at a final concentration of 400 nM), and 2×Anti-Taq Universal SYBR qPCR ProMix (Findrop) on a StepOnePlus Real-Time PCR System. The thermal cycling protocol was: 95 °C for 5 minutes; followed by 40 cycles of 95 °C for 10 seconds and 60 °C for 40 seconds; with a melting curve analysis of 65 °C for 5 seconds, and then a stepwise increase of 0.5 °C up to 95 °C. For all RT-qPCR tests, technical triplicates were conducted for each biological replicate, and calculated with the ΔΔCt method and normalized to the internal reference gene *rnpB*.

### Metadata analyses of the *Prochlorococcus*-*Uronema* co-occurrence in marine environments

To investigate the global co-occurrence patterns of *Prochlorococcus* and *Uronema*, a total of 772 datasets (metagenomes and 16S/18S rRNA gene amplicon sequences) were retrieved from the NCBI Sequence Read Archive (SRA) database, including the data from the Tara Oceans^55^, BioGEOTRACES^56^, Hawaiian Ocean Time-series^57^, Bermuda-Atlantic Time-series Study^58^, and Malaspina^59^ expeditions. To further complement these ocean expedition metagenomic datasets, keywords including “seawater” were used to search relevant literatures on PubMed, and associated metagenomic datasets were collected from the NCBI-SRA database. These datasets-associated sampling and environmental information (e.g., latitude, longitude, and water depth) were manually curated from the corresponding literatures (Table S3). These metagenomic data were processed through quality control, sequence assembly, taxonomic annotation, and abundance quantification as described (Zhou et al., 2025). Specifically, raw reads were quality-filtered using trim galore (v0.6.10)^60^ with default parameters. Clean reads from each sample were assembled individually with MEGAHIT (v1.2.9)^61^, and assembly quality was evaluated with QUAST (v5.0.2)^62^. Contigs were taxonomically annotated using both CAT (v5.2.3)^63^ and Kaiju (v1.9.2)^64^ to improve annotation accuracy and coverage. Open reading frames (ORFs) were predicted by Prodigal (v2.6.3)^65^ in meta mode and aligned against the GTDB non-redundant protein database (release_214) using DIAMOND blastp (v2.1.7)^66^. For Kaiju annotation, contigs were compared to the NCBI nr database under default settings. Finally, relative species abundances were calculated To determine the relative abundance of *Prochlorococcus* and *Uronema* in contigs, clean reads were mapped to the contigs using CoverM (v 0.6.0), with parameter ‘-contig’ and cut-off values of 75% identity and 75% alignment coverage for mapped reads, which generated coverage profiles and normalized as relative_abundance for each contig^67^. 16S and 18S ribosomal RNA gene amplicon sequencing data were collected from 90 seawater samples. To control for spatiotemporal variation and ensure the reliability of comparative analyses, samples from the same site—including data for 16S and 18S rRNA gene amplicon sequencing—were collected concurrently at an identical station during a single research cruise. Demultiplexed paired-end sequences were filtered, denoised, and merged using the DADA2 package in R (v4.1.0) to infer amplicon sequence variants (ASVs). Taxonomic assignment of prokaryotic and eukaryotic ASVs was performed using the RDP classifier against the SILVA database (v132) with an 80% confidence threshold and the Protist Ribosomal Reference database (PR2, v4.11.1), respectively^68^. Relative abundances were calculated by dividing the read count of each ASV by the total reads per sample. The Jaccard distance and Sørensen dissimilarity index were employed to assess the co-occurrence patterns between *Prochlorococcus* and *Uronema*.

### Quantification and statistical analysis

Statistical analyses were performed using one-way or two-way analysis of variance (ANOVA) for between-group comparisons, followed by Tukey’s post hoc test where appropriate. Data were fitted using both linear or nonlinear models, with the goodness of fit evaluated by the coefficient of determination (r^2^). A threshold of *p* < 0.05 was considered statistically significant. All data are presented as mean ± standard error of the mean.sss

## Reference

1. Chisholm, S.W. et al. A novel free-living prochlorophyte abundant in the oceanic euphotic zone. Nature 334, 340–343 (1988).

2. Scanlan, D.J. Physiological diversity and niche adaptation in marine Synechococcus. In Advances in Microbial Physiology 47, 1–64 (2003).

3. Hartmann, M. et al. Efficient CO_2_ fixation by surface Prochlorococcus in the Atlantic Ocean. ISME Journal 8, 2280–2289 (2014).

4. Agawin, N.S.R., Duarte, C.M., Agustí, S. Nutrient and temperature control of the contribution of picoplankton to phytoplankton biomass and production. Limnology and Oceanography 45, 591–600 (2000).

5. Schirrmeister, B.E., de Vos, J.M., Antonelli, A., Bagheri, H.C. Evolution of multicellularity coincided with increased diversification of cyanobacteria and the Great Oxidation Event. Proceedings of the National Academy of Sciences of the United States of America 110, 1791–1796 (2013).

6. Sanchez-Baracaldo, P., Bianchini, G., Wilson, J.D., Knoll, A.H. Cyanobacteria and biogeochemical cycles through Earth history. Trends in Microbiology 30, 143–157 (2022).

7. Flombaum, P. et al. Present and future global distributions of the marine Cyanobacteria Prochlorococcus and Synechococcus. Proceedings of the National Academy of Sciences of the United States of America 110, 9824–9829 (2013).

8. Partensky, F., Garczarek, L. Prochlorococcus: Advantages and limits of minimalism. Annual Review of Marine Science 2, 305–331 (2010).

9. Zubkov, M.V., Sleigh, M.A., Tarran, G.A., Burkill, P.H., Leakey, R.J.G. Picoplanktonic community structure on an Atlantic transect from 50 degrees N to 50 degrees S. Deep-Sea Research Part I Oceanographic Research Papers 45, 1339–1355 (1998).

10. Ribalet, F., Dutkiewicz, S., Monier, E., Armbrust, E.V. Future ocean warming may cause large reductions in Prochlorococcus biomass and productivity. Nature Microbiology 10, 2441–2453 (2025).

11. Zufia, J.A., Laber, C.P., Legrand, C., Lindehoff, E., Farnelid, H. Growth and mortality rates of picophytoplankton in the Baltic Sea Proper. Marine Ecology Progress Series 735, 63–76 (2024).

12. Kerkar, A.U., Sutherland, K.R., Thompson, A.W. Non-viral predators of marine picocyanobacteria. Trends in Microbiology 33, 558–568 (2025).

13. Cai, L., Li, H., Deng, J., Zhou, R., Zeng, Q. Biological interactions with Prochlorococcus: Implications for the marine carbon cycle. Trends in microbiology 32, 280–291 (2024).

14. Fenchel, T. The microbial loop-25 years later. Journal of Experimental Marine Biology and Ecology 366, 99–103 (2008).

15. Krumhardt, K.M. et al. From nutrients to fish: Impacts of mesoscale processes in a global CESM-FEISTY eddying ocean model framework. Progress in Oceanography 227, 103314 (2024).

16. Pernthaler, J. Predation on prokaryotes in the water column and its ecological implications. Nature Reviews Microbiology 3, 537–546 (2005).

17. Landry, M.R., Stukel, M.R., Selph, K.E., Goericke, R. Coexisting picoplankton experience different relative grazing pressures across an ocean productivity gradient. Proceedings of the National Academy of Sciences of the United States of America 120, e2220771120 (2023).

18. Granato, E.T., Meiller-Legrand, T.A., Foster, K.R. The evolution and ecology of bacterial warfare. Current Biology 29, 521–537 (2019).

19. Smith, W.P.J., Wucher, B.R., Nadell, C.D., Foster, K.R. Bacterial defences: Mechanisms, evolution and antimicrobial resistance. Nature Reviews Microbiology 21, 519–534 (2023).

20. Wilken, S. et al. Choanoflagellates alongside diverse uncultured predatory protists consume the abundant open-ocean cyanobacterium Prochlorococcus. Proceedings of the National Academy of Sciences of the United States of America 120, e2220771120 (2023).

21. Blasius, B., Rudolf, L., Weithoff, G., Gaedke, U., Fussmann, G.F. Long-term cyclic persistence in an experimental predator-prey system. Nature 577, 226–230 (2020).

22. Mitchell-Olds, T., Bradley, D. Genetics of Brassica rapa. Costs of disease resistance to three fungal pathogens. Evolution 50, 1859–1865 (1996).

23. Rocap, G. et al. Genome divergence in two Prochlorococcus ecotypes reflects oceanic niche differentiation. Nature 424, 1042–1047 (2003).

24. Harvell, C.D. The ecology and evolution of inducible defenses. The Quarterly review of biology 65, 323–340 (1990).

25. Magalhaes Cardoso, P.H., Balian, S.d.C., Matushima, E.R., de Padua, S.B., Martins, M.L. First report of scuticociliatosis caused by Uronema sp in ornamental reef fish imported into Brazil. Revista Brasileira De Parasitologia Veterinaria 26, 491–495 (2017).

26. Magalhaes Cardoso, P.H. et al. Scuticociliatosis caused by Uronema sp. in ten different ornamental aquarium reef fish in Brazil. Revista Brasileira De Parasitologia Veterinaria 29, e018319 (2020).

27. Grzesiuk, M., Pietrzak, B., Wacker, A., Pijanowska, J. Photosynthetic activity in both algae and cyanobacteria changes in response to cues of predation. Frontiers in Plant Science 13, 907174 (2022).

28. Connell, P.E., Ribalet, F., Armbrust, E.V., White, A., Caron, D.A. Diel oscillations in the feeding activity of heterotrophic and mixotrophic nanoplankton in the North Pacific Subtropical Gyre. Aquatic Microbial Ecology 85, 167–181 (2020).

29. Ribalet, F. et al. Light-driven synchrony of Prochlorococcus growth and mortality in the subtropical Pacific gyre. Proceedings of the National Academy of Sciences of the United States of America 112, 8008–8012 (2015).

30. Pernthaler, J. Predation on prokaryotes in the water column and its ecological implications. Nature Reviews Microbiology 3, 537–546 (2005).

31. Follett, C.L. et al. Trophic interactions with heterotrophic bacteria limit the of Prochlorococcus. Proceedings of the National Academy of Sciences of the United States of America 119, e2110993118 (2022).

32. Jover, L.F., Effler, T.C., Buchan, A., Wilhelm, S.W., Weitz, J.S. The elemental composition of virus particles: Implications for marine biogeochemical cycles. Nature Reviews Microbiology 12, 519–528 (2014).

33. Shiah, F.-K. et al. Viral shunt in tropical oligotrophic ocean. Science Advances 8, 1–7 (2022).

34. Teng, Z.-J. et al. Acrylate protects a marine bacterium from grazing by a ciliate predator. Nature Microbiology 6, 1317–1351 (2021).

35. Scanlan, D.J., Hess, W.R., Partensky, F., Newman, J., Vaulot, D. High degree of genetic variation in Prochlorococcus (Prochlorophyta) revealed by RFLP analysis. European Journal of Phycology 31, 1–9 (1996).

36. Partensky, F., Hoepffner, N., Li, W.K.W., Ulloa, O., Vaulot, D. Photoacclimation of Prochlorococcus sp. (Prochlorophyta) strains isolated from the North Atlantic and the Mediterranean Sea. Plant physiology 101, 285–296 (1993).

37. Waterbury, J., Watson, S., Valois, F., Franks, D.J.C.b.f.a. Biological and ecological characterization of the marine unicellular cyanobacterium Synechococcus. Canadian Bulletin of Fisheries and Aquatic Sciences 214, 71–120 (1986).

38. Moore, L.R. et al. Culturing the marine cyanobacterium Prochlorococcus. Limnology and Oceanography-Methods 5, 353–362 (2007).

39. Morris, J.J., Kirkegaard, R., Szul, M.J., Johnson, Z.I., Zinser, E.R. Facilitation of robust growth of Prochlorococcus colonies and dilute liquid cultures by “Helper” heterotrophic bacteria. Applied and Environmental Microbiology 74, 4530–4534 (2008).

40. Saito, M.A., Moffett, J.W., Chisholm, S.W., Waterbury, J.B. Cobalt limitation and uptake in Prochlorococcus. Limnology and Oceanography 47, 1629–1636 (2002).

41. Berube, P.M. et al. Physiology and evolution of nitrate acquisition in Prochlorococcus. ISME Journal 9, 1195–1207 (2015).

42. Rocke, E., Liu, H. Respiration, growth and grazing rates of three ciliate species in hypoxic conditions. Marine Pollution Bulletin 85, 410–417 (2014).

43. Huang, L.-F. et al. Effect of Temperature on the Growth of a Marine Heterotrophic Nanoflagellate. Acta Parasitology et Medica Entomologica Sinica 17, 10–15 (2010).

44. Li, Q., Edwards, K.F., Schvarcz, C.R., Selph, K.E., Steward, G.F. Plasticity in the grazing ecophysiology of Florenciella (Dichtyochophyceae), a mixotrophic nanoflagellate that consumes Prochlorococcus and other bacteria. Limnology and Oceanography 66, 47–60 (2021).

45. Lin, P. et al. Microplastics magnify inhibitive effects of perfluorooctanoic acid on the marine microbial loop. Environmental Research 273, 121223 (2025).

46. Liang, Y. et al. Substrate-dependent competition and cooperation relationships between Geobacter and Dehalococcoides for their organohalide respiration. ISME Communications 1, 23 (2021).

47. Ho, S.S.H., Yu, J.Z. Determination of airborne carbonyls: Comparison of a thermal desorption/GC method with the standard DNPH/HPLC method. Environmental Science & Technology 38, 862–870 (2004).

48. Saison, D., De Schutter, D.P., Delvaux, F., Delvaux, F.R. Determination of carbonyl compounds in beer by derivatisation and headspace solid-phase microextraction in combination with gas chromatography and mass spectrometry. Journal of Chromatography A 1216, 5061–5068 (2009).

49. Cantalapiedra, C.P., Hernandez-Plaza, A., Letunic, I., Bork, P., Huerta-Cepas, J. eggNOG-mapper v2: Functional annotation, orthology assignments, and domain prediction at the metagenomic scale. Molecular Biology and Evolution 38, 5825–5829 (2021).

50. Delmont, T.O., Eren, A.M. Linking pangenomes and metagenomes: The Prochlorococcus metapangenome. Peerj 6, e4375 (2018).

51. Zeng, Q., Chisholm, S.W. Marine viruses exploit their host’s two-component regulatory system in response to resource limitation. Current Biology 22, 124–128 (2012).

52. Bolger, A.M., Lohse, M., Usadel, B. Trimmomatic: A flexible trimmer for Illumina sequence data. Bioinformatics 30, 2114–2120 (2014).

53. Langmead, B., Salzberg, S.L. Fast gapped-read alignment with Bowtie 2. Nature Methods 9, 354–357 (2012).

54. Wagner, G.P., Kin, K., Lynch, V.J. Measurement of mRNA abundance using RNA-seq data: RPKM measure is inconsistent among samples. Theory in Biosciences 131, 281–285 (2012).

55. Sunagawa, S. et al. Tara Oceans: Towards global ocean ecosystems biology. Nature Reviews Microbiology 18, 428–445 (2020).

56. Anderson, R.F. Geotraces: Accelerating research on the marine biogeochemical cycles of trace elements and their isotopes. Annual Review of Marine Science 12, 49–85 (2020).

57. Karl, D.M., Church, M.J. Microbial oceanography and the Hawaii Ocean Time-series programme. Nature Reviews Microbiology 12, 699–713 (2014).

58. Michaels, A.F., Knap, A.H. Overview of the U.S. JGOFS Bermuda Atlantic Time-series study and the hydrostation S program. Deep-Sea Research Part II Topical Studies in Oceanography 43, 157–198 (1996).

59. Duarte, C.M. Seafaring in the 21st century: The Malaspina 2010 circumnavigation expedition. Limnology and Oceanography Bulletin 24, 11–14 (2015).

60. Kechin, A., Boyarskikh, U., Kel, A., Filipenko, M. cutPrimers: A new tool for accurate cutting of primers from reads of targeted next generation sequencing. Journal of Computational Biology 24, 1138–1143 (2017).

61. Li, D., Liu, C.-M., Luo, R., Sadakane, K., Lam, T.-W. MEGAHIT: An ultra-fast single-node solution for large and complex metagenomics assembly via succinct de Bruijn graph. Bioinformatics 31, 1674–1676 (2015).

62. Quast, C. et al. The SILVA ribosomal RNA gene database project: Improved data processing and web-based tools. Nucleic Acids Research 41, 590–596 (2013).

63. von Meijenfeldt, F.A.B., Arkhipova, K., Cambuy, D.D., Coutinho, F.H., Dutilh, B.E. Robust taxonomic classification of uncharted microbial sequences and bins with CAT and BAT. Genome Biology 20, 217 (2019).

64. Menzel, P., Ng, K.L., Krogh, A. Fast and sensitive taxonomic classification for metagenomics with Kaiju. Nature Communications 7, 11257 (2016).

65. Hyatt, D. et al. Prodigal: Prokaryotic gene recognition and translation initiation site identification. BMC Bioinformatics 11, 119 (2010).

66. Buchfink, B., Xie, C., Huson, D.H. Fast and sensitive protein alignment using DIAMOND. Nature Methods 12, 59–60 (2015).

67. Zhou, N. et al. Microbially-mediated halogenation and dehalogenation cycling of organohalides in the ocean. Nature Communications 16, 10670 (2025).

68. Liang, Z. et al. Unique microbiome in organic matter-polluted urban rivers. Global Change Biology 29, 391–403 (2023).

